# Bridging Imaging Users to Imaging Analysis - A community survey

**DOI:** 10.1101/2023.06.05.543701

**Authors:** Suganya Sivagurunathan, Stefania Marcotti, Carl J Nelson, Martin L Jones, David J Barry, Thomas J A Slater, Kevin W Eliceiri, Beth A Cimini

## Abstract

The “Bridging Imaging Users to Imaging Analysis” survey was conducted in 2022 by the Center for Open Bioimage Analysis (COBA), Bioimaging North America (BINA), and the Royal Microscopical Society Data Analysis in Imaging Section (RMS DAIM) to understand the needs of the imaging community. Through multi-choice and open-ended questions, the survey inquired about demographics, image analysis experiences, future needs, and suggestions on the role of tool developers and users. Participants of the survey were from diverse roles and domains of the life and physical sciences. To our knowledge, this is the first attempt to survey cross-community to bridge knowledge gaps between physical and life sciences imaging. Survey results indicate that respondents’ overarching needs are documentation, detailed tutorials on the usage of image analysis tools, user-friendly intuitive software, and better solutions for segmentation, ideally in a format tailored to their specific use cases. The tool creators suggested the users familiarize themselves with the fundamentals of image analysis, provide constant feedback, and report the issues faced during image analysis while the users would like more documentation and an emphasis on tool friendliness. Regardless of the computational experience, there is a strong preference for ‘written tutorials’ to acquire knowledge on image analysis. We also observed that the interest in having ‘office hours’ to get an expert opinion on their image analysis methods has increased over the years. In addition, the community suggests the need for a common repository for the available image analysis tools and their applications. The opinions and suggestions of the community, released here in full, will help the image analysis tool creation and education communities to design and deliver the resources accordingly.

## INTRODUCTION

Microscopy has grown tremendously in the last few decades as a discipline ranging from simple light microscopes to super-resolution and electron microscopes which can image specimens beyond the diffraction limit. In parallel, quantitative image analysis has become an integral part of microscopy. Automated microscopes now generate a large amount of data (up to TBs per day), which increasingly requires automated analysis to handle this ever-increasing data load. New modalities and submodalities of microscopy are now frequently invented, many of which require a diverse set of tools to analyze them. The requirements of the imaging community in terms of image analysis are therefore highly diverse and ever-changing.

With the aim of improving the understanding of the imaging community’s needs, the Center for Open Bioimage Analysis (COBA) along with Bioimaging North America (BINA) and the Royal Microscopical Society Data Analysis in Imaging Section (RMS DAIM) has conducted the ‘Bridging Imaging Users to Imaging Analysis’ survey in 2022. The survey consisted of 32 questions directed towards the imaging community in both life science and physical sciences on different topics such as the demographics of the participants, usage of image analysis tools, preferences for learning materials, topics of interest for future workshops, image analysis experiences, suggestions for image analysis tool creators and users. The responses received were compared with the results of the 2020 Bioimage analysis survey conducted by COBA to understand the preferences and needs of the imaging community and thereby develop and disseminate the resources accordingly.

## RESULTS

### Participants, their work type, and computational skills

The survey was conducted from May 2022 to July 2022; the survey was open to the general public and promoted in the imaging community through the Images2Knowledge (I2K) and Electron Microscopy and Analysis Group (EMAG) conferences, the image.sc forum^1^, Microforum, Twitter, Confocal, ImageJ, and BioImaging North America (BINA) listservs. In contrast to a previous survey from this team^2^, which was limited to bioimage analysis only, questions were added around physical science analysis as well. The final results contain 493 participants from a variety of roles and scientific experiences (Fig 1A&B). The most common training category is ‘Cell/Molecular Biology’. While our experience and data such as CellProfiler website analytics (Fig S1B) indicate global interest in accessing image analysis, most survey participants were from Europe and North America (Fig S1A) possibly because of the medium of distribution of the survey. The geographic results, along with the fact that students and postdoctoral fellows together only make up less than 40% of our sample, mean our results if left unfiltered do not create an unbiased sample of the imaging community as a whole; nevertheless, with cautious examination and subsampling, trends and conclusions can be drawn.

**Figure 1 -.**
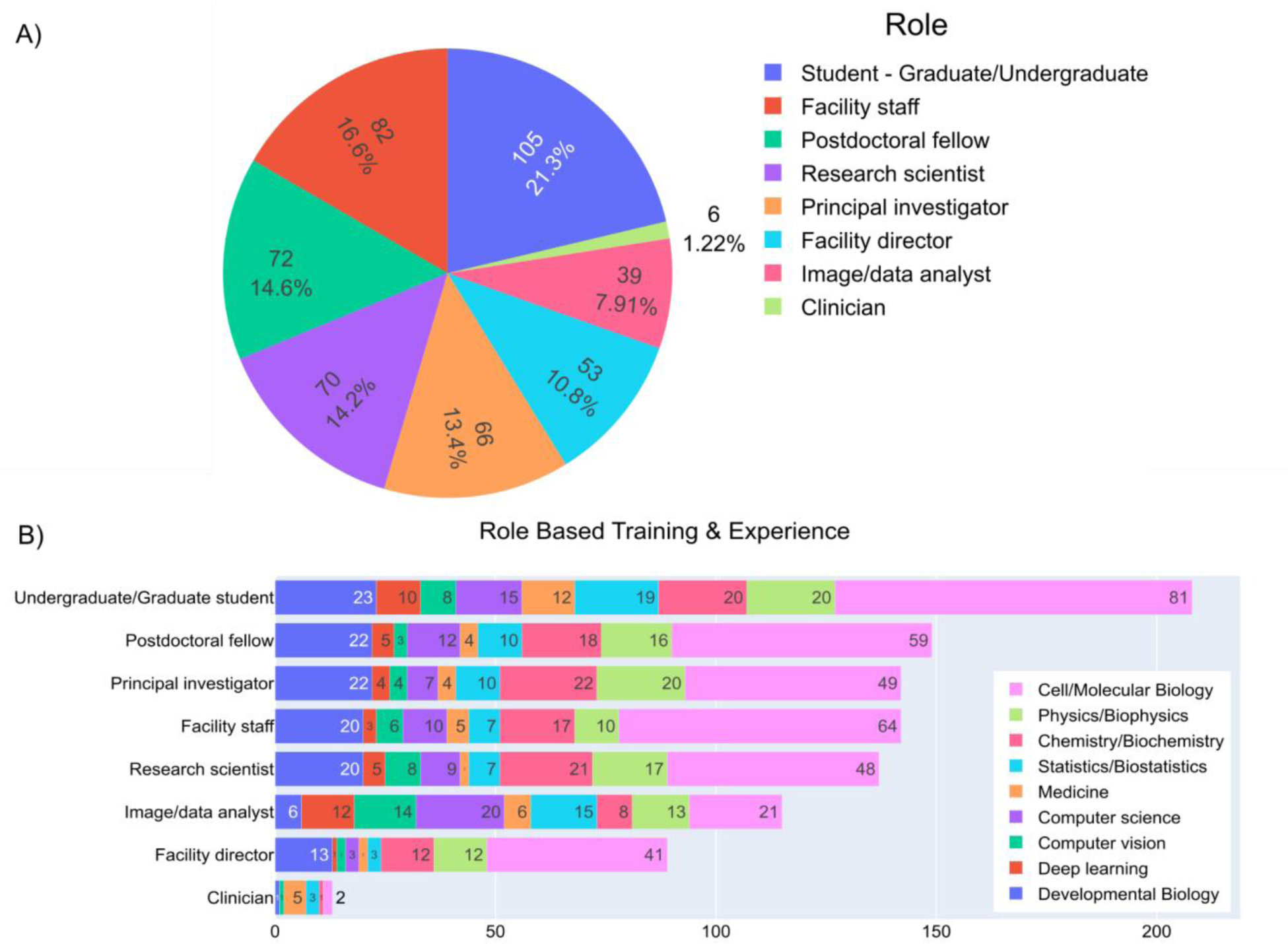
Survey respondents’ roles and training histories vary across the sampled responses. A) Answers to the multiple-choice question “Which of the following roles best describes you?”. B) Answers to the check-box question “Which of the following do you have significant formal training in or experience with? Select all that apply.” Responses were categorized based on the answers provided for part A.

Five major descriptors were used to break down other survey respondents: choice of life science vs physical science questions, self-reported computational skill level, self-reported computational comfort level, primary work classification (between Imaging, Analyst, or Balanced), and trainee status; details can be found in the methods section (Fig S2A-C). A majority of the respondents were under the ‘Balanced’ work type irrespective of the domain and trainee status (Fig 2, S2D-G); specialists in either imaging or analysis were more common in non-trainee roles, possibly due to the >25% of respondents who describe themselves as facility directors or staff. Most of the participants had moderate computational skills except for the ‘Analysts’ in life sciences, whose self-reported computational skills were higher than the other work types. Respondents were also asked to describe their ‘Comfort in developing new computational skills’; as one might expect, more computational job roles and self-reported skill levels are associated with higher comfort. (Fig 2). Overall, most of the participants of the survey including the trainees and non-trainees had moderate computational skills in both life and physical sciences (Fig 2, S2E-G).

**Figure 2 -.**
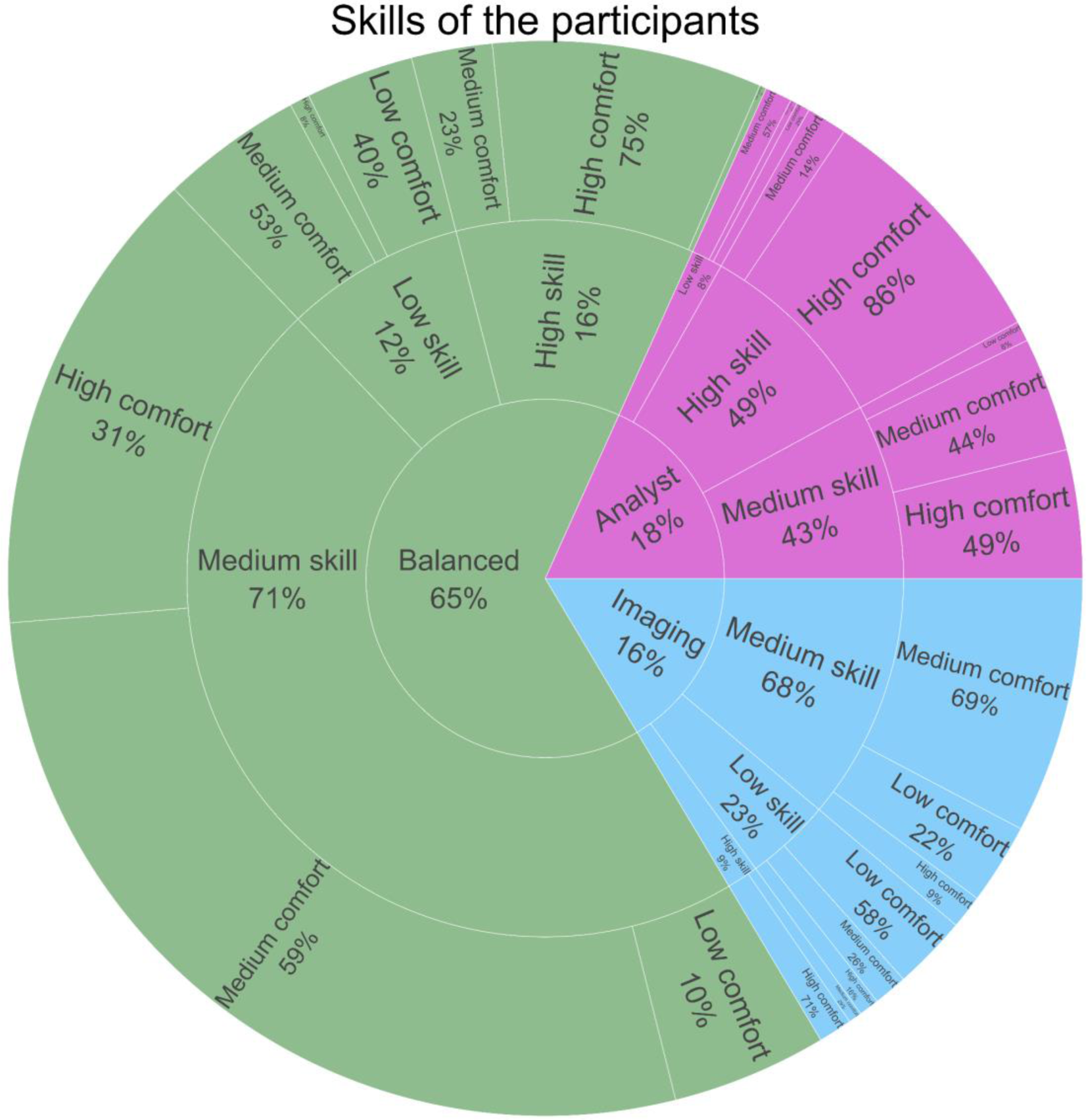
Skills of the participants. Breakdown of answers to the multiple-choice questions “How would you describe your work?”, “How would you rate your computational skills?” and “How would you rate your comfort in developing new computational skills?” Percentages were rounded to the nearest percent; in outer wedges, percentages are of the adjacent inner wedge population. See methods and Supplemental Figure 2 for fuller descriptions of each category; interactive versions of these plots are available at https://broad.io/2022SurveyApp.

### Images and image analysis tools

Understanding the kinds of images most commonly analyzed by the imaging community is important to help developers to design their tools accordingly. When surveyed, the majority of life science participants wanted to analyze fluorescent images that were manually acquired followed by the ones that were acquired in an automated manner (such as by a plate scanning microscope) (Fig 3A). Fluorescence microscopic images were also the most analyzed images in the physical sciences, but physical science respondents were far more likely to be doing electron microscopy than life scientists. (Fig 3B). 2D images were the most commonly generated images, followed by 2D +time, 3D, 3D+ time, and large-volume 3D images (Fig 3A&B).

**Figure 3 -.**
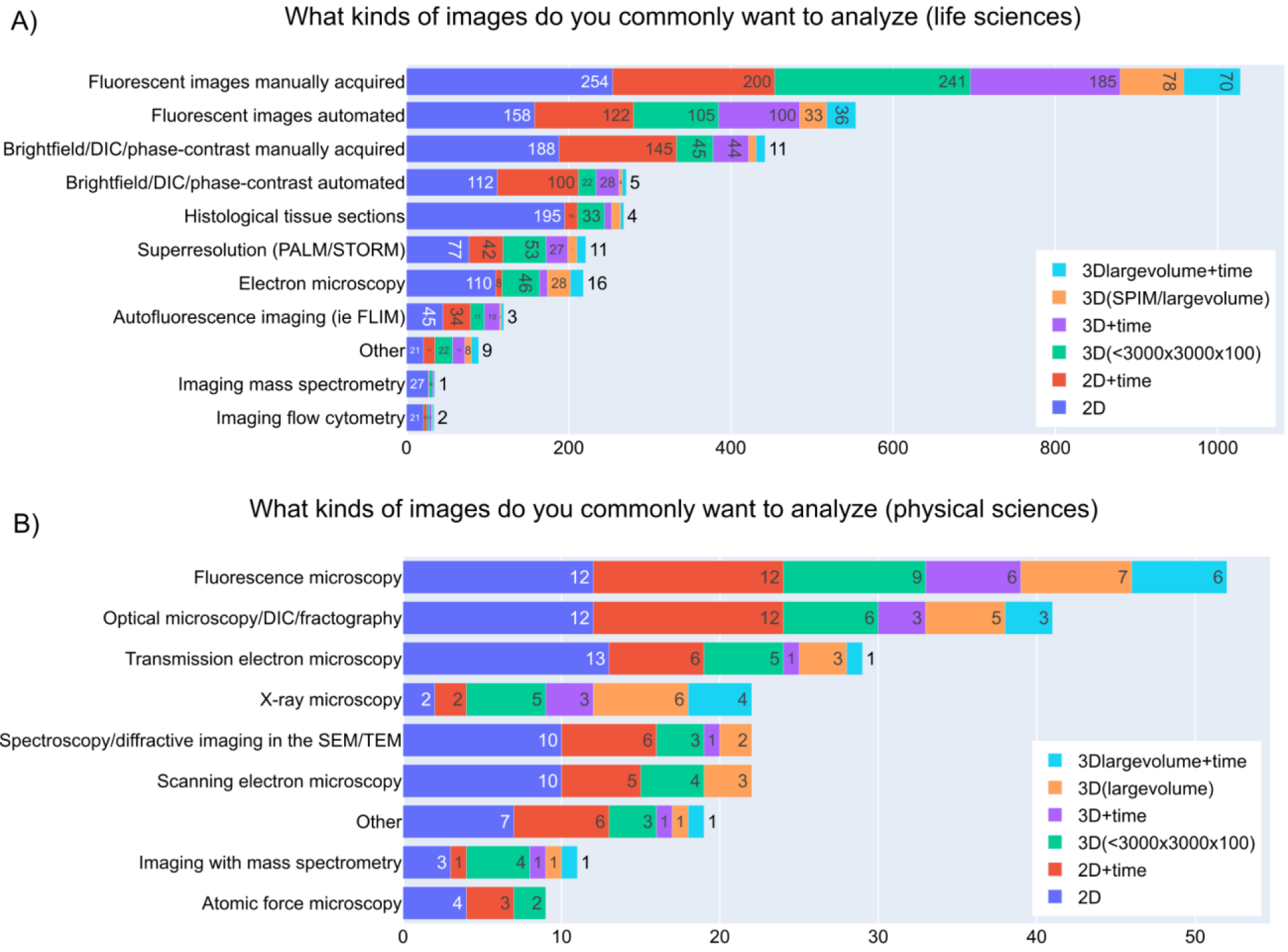
Kinds of images analyzed. A) Answers to the checkbox grid question “What kinds of images do you commonly want to analyze (select all that apply)?” under the “Life Sciences Image Analysis” section. B) Answers to the checkbox grid question “What kinds of images do you commonly want to analyze (select all that apply)?” under the “Physical Sciences Image Analysis” section.

Participants were next asked about the image analysis tools that they use; as in our previous survey^2^, we observed open-source point-and-click software are the most used image analysis tools (Fig 4), which we hypothesize may be due to any or all of the following - ease of use, ease of access, availability of tutorials, or the ability to perform analyses without prior programming knowledge. No-code-required tools are not necessarily needed by every member of the community with approximately ¾ of participants reporting using computational libraries and scripts for analyzing their images at least sometimes; while sampling bias no doubt partially contributes to this number, it does indicate a somewhat higher community competency with scripting than self-reported comfort levels might suggest. (Fig 4 A&D). The frequency at which the scripts are used is higher among the physical science participants (Fig 4 C&F, S3), which likely relates to the higher-reported computational skills of physical science vs life science participants (Fig S2 F&G). The higher-reported computational skills of physical science participants might be reflective of a higher tendency of undergraduate programs in these disciplines to include programming. The techniques available in physical sciences imaging (electron microscopy, AFM, spectrum imaging etc.) may also have fewer established point-and-click software packages for analysis, particularly in some specialized techniques.

**Figure 4 -.**
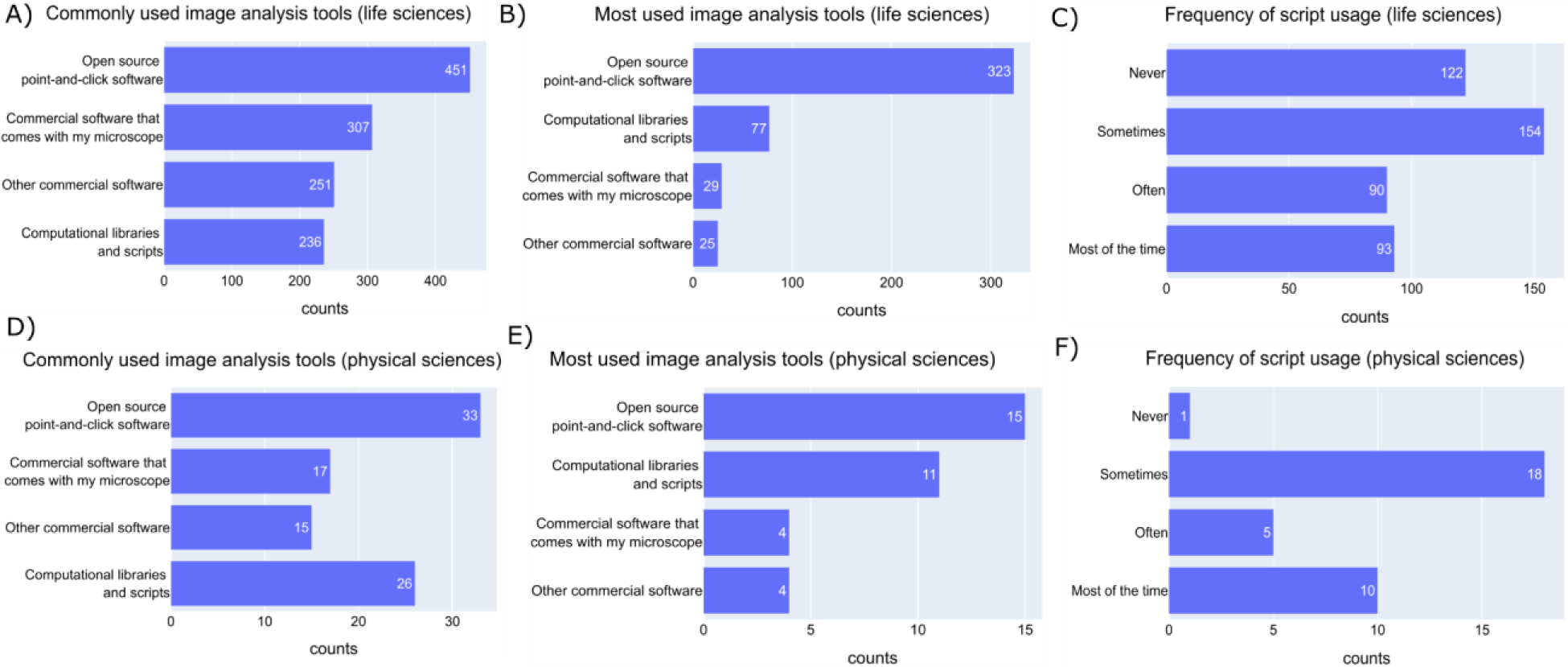
The community prefers open-source point-and-click software. A) Answers to the multiple-choice question “What image analysis tools have you used before? (check all that apply)” under the “Life Sciences Image Analysis” section. B) Answers to the checkbox question “What image analysis tools do you use the most?” under the “Life Sciences Image Analysis” section. C) Answers to the question “How frequently do you use scripting to solve image analysis problems?” by ‘Life Science’ participants. D) Answers to the multiple-choice question “What image analysis tools have you used before? (check all that apply)” under the “Physical Sciences Image Analysis” section. E) Answers to the checkbox question “What image analysis tools do you use the most?” under the “Physical Sciences Image Analysis” section F) Answers to the question “How frequently do you use scripting to solve image analysis problems?” by ‘Physical Science’ participants.

### Solving image analysis problems

Given the complexity of images often generated, it is common for participants to move beyond simple analysis using single methodologies. When the participants were asked about the approaches they use, it was clear that they prefer to use the tools that they have already used and are comfortable with. Participants rely on the internet, scientific literature, and their colleagues’ protocols for any problems they face when coming up with a solution for analyzing their images. Despite the creation of the Scientific Community image forum^1^ as a central hub for questions around image analysis and with answers provided by the experts in the field, usage remains comparable to 2020 levels^2^ (Fig 5A) and low (28%) even among analysts (Fig S4A).

**Figure 5 -.**
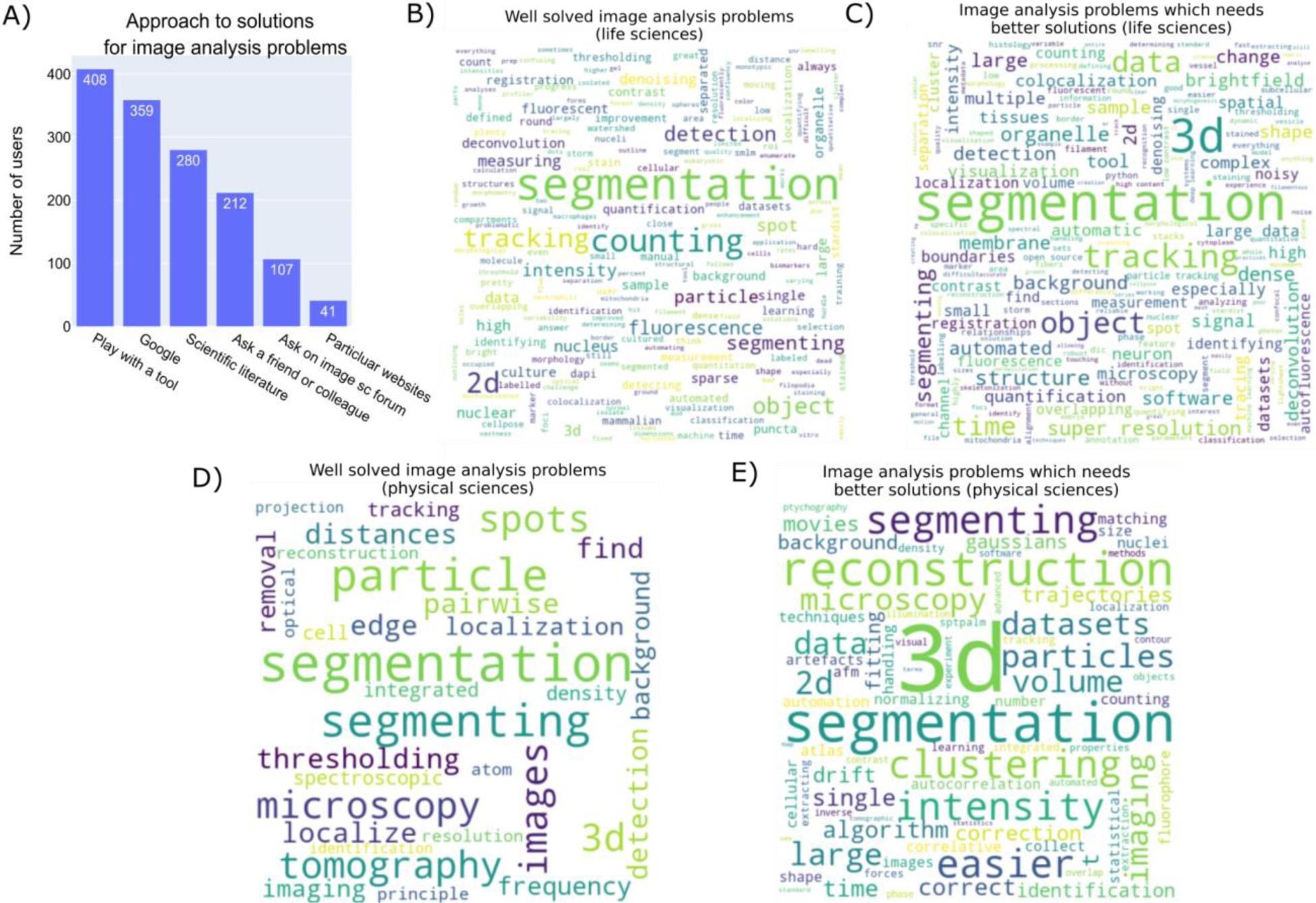
Solving image analysis problems. A) Answers to the checkbox question “How do you generally go about solving an image analysis problem? Check the approach(es) you use the most.” B) Wordcloud representation of the unigrams of the answers by ‘life science’ participants to an open-ended question “What image analysis problems (i.e. finding nuclei, tissue analysis, analysis of super-resolution data, etc) do you think are generally well-solved?” C) Wordcloud representation of the unigrams of the answers by ‘life science’ participants to an open-ended question “What image analysis problems (i.e. finding nuclei, tissue analysis, analysis of super-resolution data, etc) do you wish had easier/better solutions?” D) Wordcloud representation of the unigrams of the answers by ‘physical science’ participants to an open-ended question “What image analysis problems (i.e. finding nuclei, tissue analysis, analysis of super-resolution data, etc) do you think are generally well-solved?” E) Wordcloud representation of the unigrams of the answers by ‘’physical science’ participants to an open-ended question “What image analysis problems (i.e. finding nuclei, tissue analysis, analysis of super-resolution data, etc) do you wish had easier/better solutions?”

We also asked open-ended questions about what the participants thought were the well-solved image analysis problems and also about the problems that need better solutions. Respondents considered ‘segmentation’ to be an image-analysis problem that is both well-solved and needs a better solution, reflective of both segmentation’s centrality to many image analysis problems as well as how wide the variety of segmentation problems are (Fig 5B-E). Three-dimensional image analysis and tracking are listed as major needs, similar to the answers that we received from the 2020 image analysis survey participants (Fig S4B&C). A notable difference between life and physical sciences were major topics in analysis needs, with life science respondents clearly highlighting tracking as a major issue (with the assumption this relates to cell tracking) and physical science respondents highlighting reconstruction (although it’s unclear whether this corresponds to 3D reconstruction in tomography or reconstruction of object phase in electron microscopy techniques).

### Experience in image analysis

To get a general idea of the participants’ experience in imaging and image analysis, we asked whether they have attended any workshops/conference sessions/conferences specific to these areas. The responses indicate that the participants have had considerable exposure to the field of imaging and image analysis as most of the participants have attended a workshop/tutorial on these topics; specifically for analysis, approximately ¾ of respondents had attended or considered attending at least one image analysis event, though only about ¼ described attending “some” or “many” (Fig 6 A&B). Workshops like NEUBIAS have been consistently quoted by the community members as one of the most useful workshops showing the interest of the imaging community in learning about the advancements in the field (Fig 6C)^2^. The participants found these experiences useful for the following reasons - the workshops/conferences provided hands-on experience in working with many tools, interaction with the experts in the field, detailed theory, easy-to-follow video tutorials, and Q&A sessions.

**Figure 6 -.**
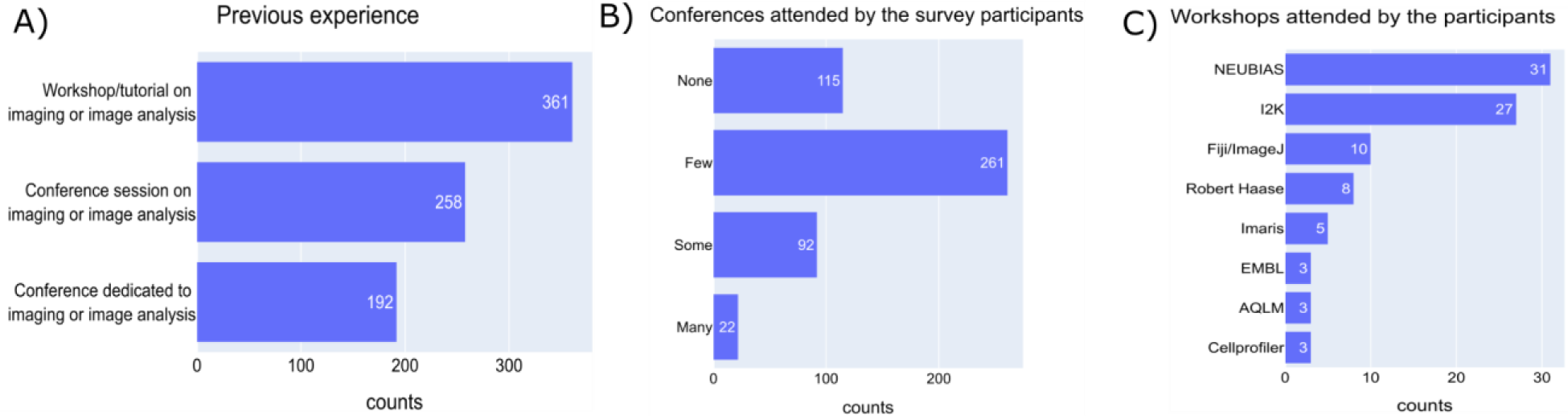
Experience in image analysis. A) Answers to a multiple-choice question “Please select any of the following you have attended in the past” B) Answers to the checkbox question “Are there any image analysis workshops, tutorials, or conferences you are aware of and attended or considered attending? If so, how many?” C) Answers to an open-ended question “Are there any image analysis workshops, tutorials, or conferences that you have participated in and found particularly helpful? If yes, what made them beneficial?”

### Topics of interest and preferable methods for learning image analysis

Given a choice, the community prefers to learn image analysis practices that are more specific to a certain sub-discipline, the methods to analyze large images, and tools to visualize the results, with more than half of participants describing themselves as “very interested” (Fig 7A). The strong preference for sub-discipline specific learning is consistent with previous results^2^; while it is unsurprising that most users want the tools only most relevant to them, the extra time required to tailor a generalist tool (and/or its training materials) to a specific audience often must be balanced against other aspects such as bug fixes and feature additions.

**Figure 7 -.**
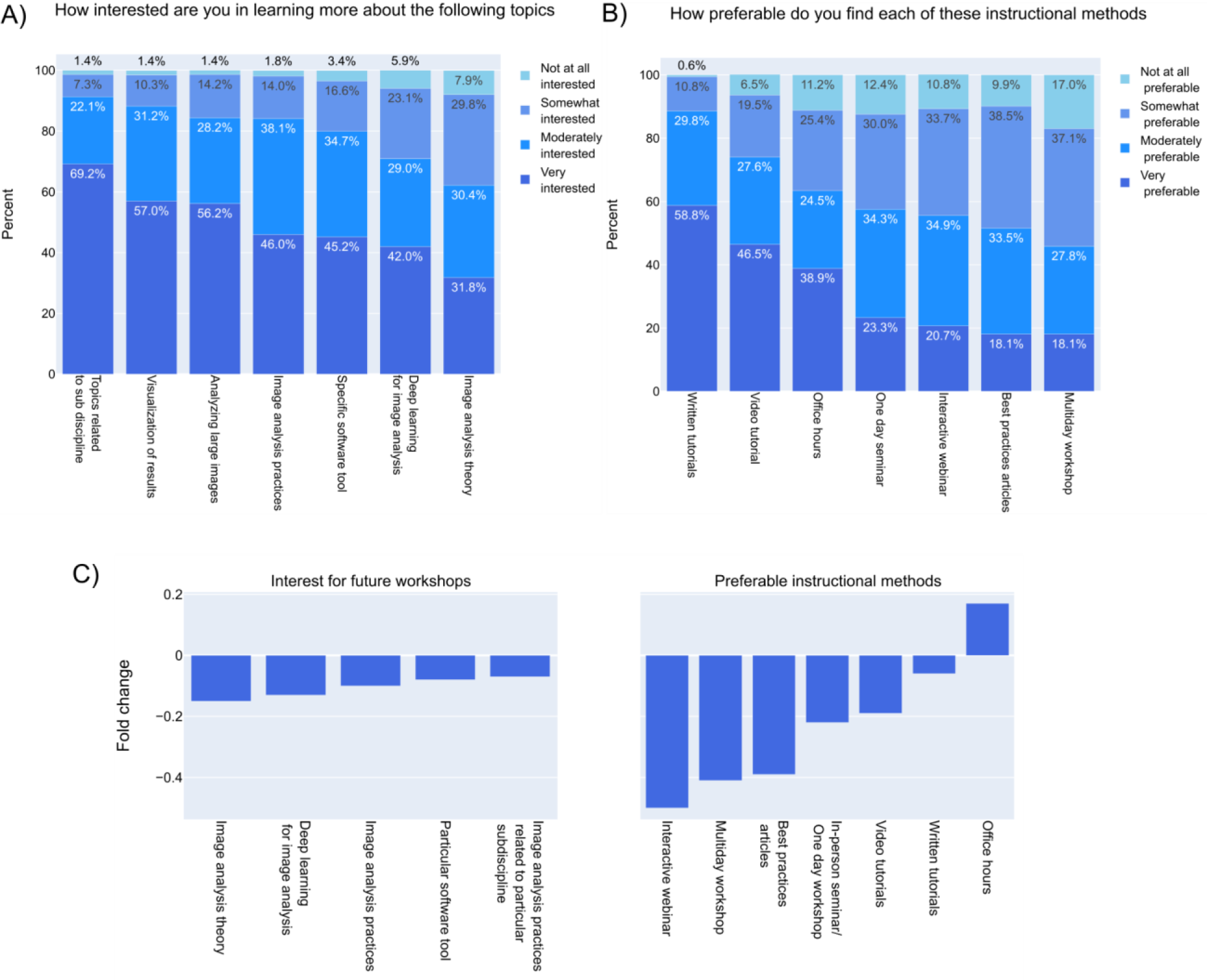
The imaging community prefers to learn about customized image analysis tools at their own pace. A) Answers to a multiple-choice grid question “How interested are you in learning more about the following topics?” B) Answers to a multiple-choice grid question “In regards to learning more about image analysis, how preferable do you find each of these instructional methods?” C) Percent change in the “Very interested/Very preferable” category of part A and B compared to the results from the 2020 bioimage analysis survey.

Based on the comments given by the participants on what made the conferences/workshops on image analysis very beneficial, it is clear that modes of instruction have played a significant role. The participants were asked about the modes of delivering knowledge on image analysis to be aware of their preferences. Written tutorials have always been the highly favored method to acquire knowledge on image analysis (Fig 7B)^2^. The main advantages could be self-paced learning, step-by-step instructions which the users could follow more easily, and flexibility in usage. To know if this is true across people with different computational skills and work types, the preferable instructional methods were cross-matched with specific categories. Regardless of the computational knowledge and work type, ‘written tutorials’ are highly preferred by the imaging community (Fig S6). Participants who fell under the category of ‘low computational skills’ and ‘imaging’ work type prefer video tutorials and office hours along with written tutorials (Fig S6). The imaging users may not possess comprehensive awareness of the availability of resources that are required for their image analysis. In such instances, expert guidance and video tutorials would be more efficient than written tutorials alone. It is noteworthy that in contrast to all other topics or instructional methods, interest in office hours increased since 2022 (Fig 7C)^2^, which could be because of the community’s general interest in learning about customized methods that would work for their own images and also for expert guidance. Interactive webinars had the largest decrease in interest between 2020 and 2022, possibly because they were so heavily leaned on as primary instructional methods in 2020 and 2021 when COVID restrictions made other instructional methods far less common; if interest continues to decline in future years, instructors may need to reassess such formats in terms of desirability and effectiveness.

### Suggestions for future workshops

Having asked about the topics and preferable methods for delivering knowledge on image analysis, the participants were next asked about the conferences that would benefit from such image analysis sessions. The respondents proposed that including image analysis workshops/sessions would be helpful for the attendees in conferences such as the American Society for Cell Biology (ASCB), Microscience Microscopy Congress (MMC), European Light Microscopy Initiative (ELMI), Association for Biomolecular Resource Facilities (ABRF), Focus on Microscopy (FOM), Biophysical Society and developmental biology conferences (Fig 8A). Some of these meetings have started introducing such sessions in recent years (ASCB, MMC, ELMI, FOM) and offering satellite workshops on image analysis topics; however, some of the sessions were more oriented towards tool highlighting than workflow construction.

**Figure 8 -.**
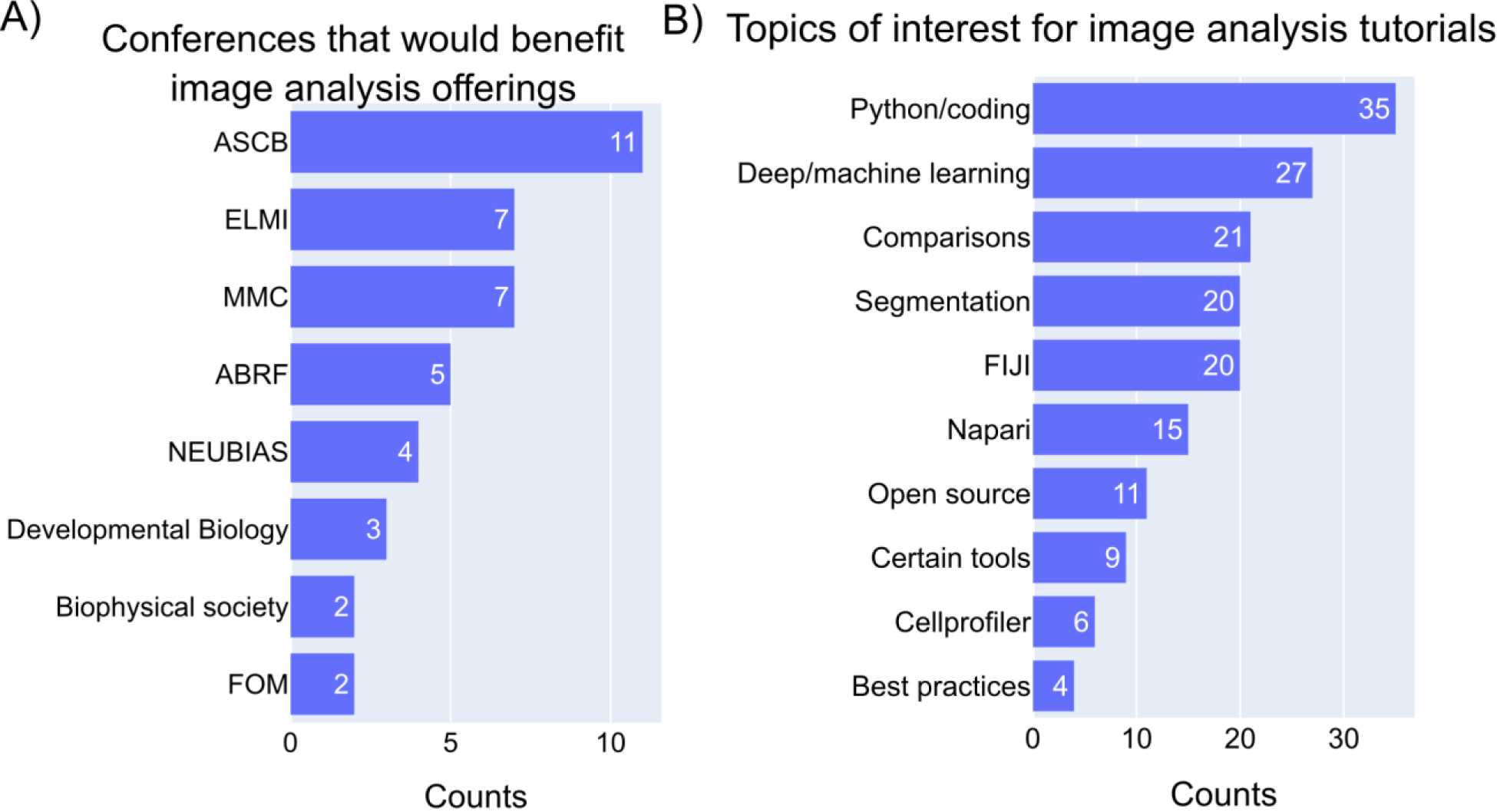
Topics of interest for the image analysis sessions in a conference. A) Answers to an open-ended question “Are there any image analysis workshops, tutorials, or conferences that you have participated in and found particularly helpful? If yes, what made them beneficial?” Unigrams and bigrams were counted from the answers and the meaningful words were plotted. B) Answers to an open-ended question “What specific topics (i.e. overviews of a particular tool, comparisons between pieces of software, or how to use a certain tool for a certain kind of experiment) would you like to see prioritized for future image analysis workshop and tutorial offerings?” Unigrams and bigrams were counted from the answers and the meaningful words were plotted.

In regards to the content of the workshops/tutorials on image analysis, the participants suggested including the methods of using Python (coding/scripting), and deep or machine learning for image analysis, the comparisons of different image analysis tools, segmentation methods, ImageJ/Fiji plugins and macros, open-source software, certain tools of interest, CellProfiler, and also the best practices in image analysis (Fig 8B). Advancements in imaging technologies have made image analysis a multi-disciplinary field and the imaging community’s curiosity to learn coding/scripting/machine learning in image analysis is reasonable. Getting comfortable with computational skills gives the end-user an opportunity to automate the image processing steps and helps in analyzing difficult-to-analyze images.

### General suggestions

Participants were asked to provide suggestions on the roles of tool creators as well as tool users in improving image analysis; the responses were then categorized based on the work type to understand the opinions of each group, since each group may have particular insight into their own role as well as the roles of others. In response to how tool creators could improve image analysis, regardless of work type, the common suggestions were the need for documentation, open-source software, video tutorials on how to use the software, user-friendly intuitive GUI, help with installation, and example data to practice the software. The need for video tutorials is highly quoted by the ‘imaging’ participants when compared to the other two groups. There was also a suggestion on recognizing the contribution of the developer through awards or incentivizing the developers who are creating user-friendly codes (Fig 9A).

**Figure 9 -.**
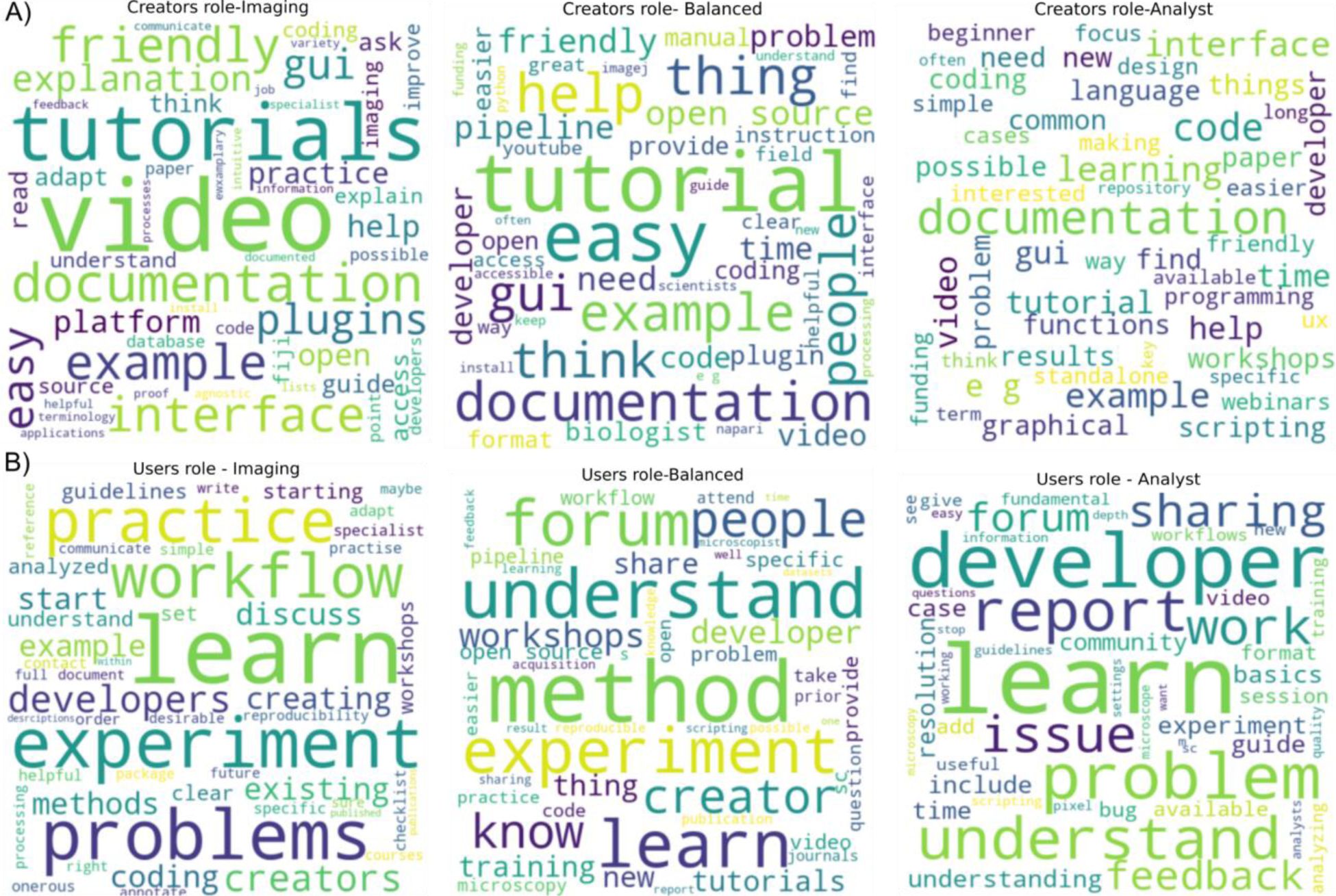
More documentation and feedback are necessary. A) Answers to an open-ended question “What do you think analysis tool CREATORS (such as software developers)could/should do to make image analysis better and more successful? How best could we encourage them to do it?” was categorized based on the “work type” as described in supplementary S2A and the unigrams of the answers are represented as wordclouds. Word clouds were made with a cut-off of 50 words. B) Answers to an open-ended question “What do you think analysis tool USERS (such as microscopists) could/should do to make image analysis better and more successful? How best could we encourage them to do it?” was categorized based on the “work type” as described in supplementary S2A and the unigrams of the answers are represented as wordclouds. Wordclouds were made with a cut-off of 50 words.

On the suggestion of users’ role to make image analysis better, **‘**Imaging’ participants feel that the users should practice the image analysis workflows with the examples, discuss the problems with developers and learn the basics by attending workshops. ‘Balanced’ and ‘Analyst’ work types suggest the users to understand fundamental image analysis by attending workshops/conferences/basic courses, learning the basics of coding/scripting, reporting image analysis problems with the developer through forums, sharing videos of how-to tutorials from users’ perspective, discussing the microscopy experiments with an image analyst before image acquisition, and providing feedback to the developers (Fig 9B). In general, frequent, and constant interaction between the tool user and developer is a common suggestion.

### Any other thoughts

A final, open-ended question invited respondents to express “any other thoughts” that they had. The respondents expressed their needs for more user-friendly tools/plugins, a common repository for the available image analysis tools with their applications, and guides/checklists for both the users and the developers. There were also requests for detailed image analysis pipelines, training the microscope core facility staff, and also to include more techniques/tools for analyzing histology images and for the images generated using other modalities such as multi-photon microscopy and scanning probe microscopy. Many of these requests were also received from the 2020 survey participants, indicating the importance of these issues^2^.

## DISCUSSION

We have reported the responses provided by 493 participants regarding their needs and suggestions for image analysis, which were collected through a survey questionnaire containing 32 questions. Combining input from physical and biological sciences was added to the survey this year, in order to determine consistent image analysis needs across the two communities. Our data from the survey will be helpful for the tool creators to design the tools based on the suggested needs and also help COBA in designing and providing the resources accordingly.

Our data suggest a significant undersampling of trainees, physical scientists, users exclusively involved in imaging, and respondents outside of North America and Europe.The undersampling of physical sciences data makes drawing firm comparisons difficult, and future surveys will aim to target this community further. These undersamplings emphasize the importance of looking beyond overall responses and tailoring analyses to particular subgroups to try to more accurately reflect the perspectives of the imaging community. Involving imaging core facilities at various universities and imaging consortiums that are focused on other continents such as the African Bioimaging Consortiums would broaden the representation of the imaging community in future such surveys. Incorporating input and feedback from a broader community will be helpful in better understanding the users’ needs allowing the tool developers to prioritize the efforts accordingly.

When approached as a whole, some inherent contradictions can be found within our survey results: a major one is that users and developers are overwhelmed with the variety of new tools being created, such that they feel more central repositories are needed, but users simultaneously request more emphasis on field-specific rather than general approaches and developers in response create new tools rather than adding functionalities to existing ones. These contradictions are understandable, especially in the light of competing incentive structures: users want to focus on image analysis more as a tool than as a discipline when they are trying to publish scientific results in a time and/or funding-constrained manner, so naturally want something targeted; developers may find it easier to publish and/or receive funding for a new tool than for maintenance and feature addition for existing tools. While solving this contradiction is beyond this group’s power, it should be noted that some repositories such as the BioImage Informatics Index (BIII, https://biii.eu/)3 and the Workflow Hub (https://workflowhub.eu/) do already exist. Moreover, resources collating conference and training materials are available, such as MicroscopyDB (https://microscopydb.io/), maintained by Global Bioimaging and BINA, and community-led efforts such as FocalPlane (https://focalplane.biologists.com/). To better serve community needs, forums can include ‘resources’ pages with pointers to these centralized sources alongside specific updates on the latest resources and their applications. Some centralized repositories such as BIII allow for field and/or function-specific tagging, which end users wishing to help their communities could prioritize adding for their sub-discipline; other repositories could consider adding such tags.

Another point that stands out when analyzing the differences between end-users’ and tool developers’ needs and requests relates to the need for more communication between the two communities. The authors feel that this gap is often bridged by the figure of the image analyst, who can offer dedicated support to the end users, while also being able to provide detailed feedback to the developers. Image analysts are often associated with imaging facilities, however, more and more examples of dedicated centers and consortia are reported. One example is AI4Life (https://ai4life.eurobioimaging.eu/), a Horizon Europe-funded consortium offering research services and infrastructure to support life scientists in the adoption of machine learning solutions for image data. In Germany, a centralized initiative (NFDI4BIOIMAGE, https://nfdi4bioimage.de/en/start/) has recently kicked-off to provide scientists from all natural science and biomedical research fields with workable and trusted solutions to handle image data. Moreover, research infrastructures offering open access to imaging technologies, such as Euro-Bioimaging (https://www.eurobioimaging.eu/), have started offering image and data analysis services for biological and medical data. It is likely that such projects will have a positive impact in improving the communication between end-users and developers, by providing the role of the image analyst who can communicate with both and facilitate each group’s interactions.

The Scientific Community image forum^1^ was introduced as a central platform to discuss, share, and provide help on the issues faced when acquiring or analyzing images. While overall traffic levels imply that utilization of the Image sc forum has steadily increased over the years, the number of survey participants saying it is among the approaches they use the most did not increase here, nor was it >30% in any subgroup including analysts. To address this, future efforts can be made to bring awareness about the existence and benefits of the forum through image analysis workshops/tutorials (ideally not ones that users must seek out, but ones attached to discipline-specific conferences) and also through targeted education opportunities for imaging facility staff, who can then relay information about best practices and about where to ask further questions to their users. Use of the forum is one of the most direct ways for an imaging tool user to gain access to expert help in their image analysis, as it is essentially a 24/7 virtual office hour; beyond direct help, creating a public record of problems encountered and solved makes it easier for the next user with the same issue to overcome the challenges they are facing. Questions of all difficulty levels but especially from beginners are warmly encouraged; users who are uncomfortable attaching their real names to their posts are welcome to create pseudonymous accounts.

Beyond documentation of individual issues encountered, broader training materials on how to solve particular problems (ideally created in such a way that people with any/no computational background can follow them) are clearly still needed in greater numbers as well as tailored to more individual communities such as multi-photon microscopy and scanning probe microscopy. Broader still were the requests from the imaging community, irrespective of the work type, to have a checklist or guidelines for imaging and image analysis; a number of such works are under construction in a variety of consortia to help users in planning their microscopy experiments, analyzing, and publishing images^4, 5^.

One of the most surprising results in our opinion was the decrease in interest in deep learning. There was an approximately 2-fold increase in the peer-reviewed papers published with the terms ‘deep/machine learning/artificial intelligence’ (Fig S8) during the period of 2020 to 2022, reflecting the overall general increasing interest in creating machine learning models or applying them in scientific discovery. This is reflected in several new (or “newly-deep”) deep learning/machine-learning based tools designed to be easy for end users, such as Cellpose^6^, StarDist^7^, ilastik^8^, DeepImageJ ^9^ and Piximi^10^, that were introduced for image analysis during the last few years. While overall trends were towards fewer “very interested” responses in 2022 vs 2020, deep learning represented the second largest decrease in topic interest. While we cannot rule out that people were less “very interested” in learning about deep learning because increasing fractions of users mastered it in that two year window, we think a more likely explanation is that greater education and outreach, as well as continued improvements in installation and access, will be needed before end users trust these tools enough to become excited about applying them to their own work. Deep learning tools are of course not always the best tool for a job, especially when large amounts of training data are unavailable and/or painful to make; improved human-in-the-loop ^11^ approaches to reduce the annotation burden and improved self-supervised training approaches may reduce this burden in the future. Finally, it is possible that the reduced uptake of such methods might relate to the lack of suitable local infrastructure and training data. Some efforts in democratizing access via Google Colab (ZeroCostDL4Mic^12^, https://henriqueslab.github.io/resources/ZeroCostDL4Mic/) and creating hubs for bringing AI models to researchers (Bioimage Model Zoo, https://bioimage.io/) have been made in recent years, and it will be interesting to see if they will have an impact in the near future.

## CONCLUSION

While the last several years have brought significant advances in many subfields of microscopy analysis, these results suggest that we as a community still have much work to do in order to empower and/or encourage imaging users to fully embrace analytical approaches and tools. We hope that these results, as well as the analysis of what has and has not changed since our initial report, can help further motivate researchers to “cross the divide” as well as motivate journals, universities, and funding agencies to assess how they can help in this process, such as by increasing access to statistical and computational training for students and early career researchers or by funding developers to create more friendly and more tailored instructional material. Unlike in our past survey, this paper provides the underlying response data in fully intact form, which we hope will encourage stakeholders to perform their own subsetting and analysis of their own section of the community.

Ultimately, the development of image analysis tools, workflows, and approaches is an iterative process requiring input from all stakeholders. No single group or community in isolation can accomplish these goals on their own; all must work together in order to fully overcome the past history of microscopy as a purely quantitative science. We hope this data adds to previous work in order to help light the way.

## PRACTITIONER POINTS

1. Gaps remain between image analysis tools developers and tool users; image analysts as a professional class may help close this gap. The Scientific Community Image Forum (forum.image.sc) is proposed as a central organizing platform.
2. Major requests for developers included increased video tutorials and office hours.
3. Tailoring of tools to particular subfields as well as including image analysis into topic-specific conferences may help increase image analysis tool adoption.

## METHODS

“Bridging Imaging Users to Imaging Analysis” survey conducted by the Center for Open Bioimage Analysis (COBA), Bioimaging North America (BINA), and the Royal Microscopical Society (RMS) was made available in Google forms to the imaging community through the Images2Knowledge (I2K) and Electron Microscopy and Analysis Group (EMAG) conferences, the image.sc forum^1^, Microforum, Twitter, Confocal, ImageJ, and BioImaging North America (BINA) listservs. The responses of the survey were exported to tables, analyzed, and graphed in Jupyter Notebook (6.4.12)^13^ using Python3(3.9.13)^14^, pandas(1.4.4)^15^, plotly(5.9.0)^16^, matplotlib(3.5.2)^17^, numpy(1.24.2)^18^, wordcloud(1.8.2.2)^19^, kaleido (0.1.0.post1) and sci-kit learn(1.0.2) ^20^ libraries. Interactive versions were made with Streamlit^21^.

Participants who answered as ‘Undergraduate/Graduate student’ and ‘Postdoctoral fellow’ to the question ‘Which of the following roles best describes you?’ were categorized as ‘Trainees’ and the rest of the roles were categorized as ‘Nontrainees’. Answers to the questions on ‘work description’, ‘computational skills’, and ‘comfort in developing new computational skills’ were grouped into three categories. Scale values of 1 & 2 were grouped as ‘Imaging/Low skill/Low comfort’, values of 3 to 5 as ‘Balanced/Medium skill/Medium comfort’, and values 6&7 as ‘Analysts/High skill/High comfort’ based on the relevant questions.

For the fold change in the interest level or the preferable methods, percentage fold change was calculated with responses received under the category ‘Very interested’/ ‘Very preferable’ for the questions - ‘In regards to learning more about image analysis, how preferable do you find each of these instructional methods?’ and ‘How interested are you in learning more about the following topics?’ from the 2020 and 2022 surveys.

The open-ended questions on solving image analysis problems, creators and users role were analyzed using wordcloud library where the answers were split into single words and the frequency of occurrence of the words decided the font size of the words in the wordcloud plot. Words that appeared in the related questions and the words that were not providing useful information were removed from the wordcloud plot along with the standard ‘English’ stopwords available in the wordcloud library. Unigrams, bigrams, and trigrams were generated from the responses to the open-ended questions ‘Are there any conferences you’ve attended in the past that you think would particularly benefit from the addition/expansion of image analysis offerings?’ and ‘What specific topics (i.e. overviews of a particular tool, comparisons between pieces of software, or how to use a certain tool for a certain kind of experiment) would you like to see prioritized for future image analysis workshop and tutorial offerings?’ using CountVectorizer library. The words/ conferences that were making it to the top of the list and also the meaningful words were counted and plotted as a graph.

## ACKNOWLEDGMENTS

The authors gratefully acknowledge members of the Cimini and Eliceiri labs for feedback on the survey questions. The authors thank Erin Weisbart, Mario Cruz, Ellen Dobson, Rebecca Senft for their suggestions and Rebecca L. Ledford for help on the data statement and publicizing the survey. They also acknowledge the strong support of Vanessa Orr, Nikki Bialy and the Image Informatics working group of BioImaging North America (BINA) in giving useful feedback and promoting the survey. They are additionally grateful to the 493 unnamed participants, without whom this work would have of course not been possible.

## DATA AVAILABILITY STATEMENT

The survey questions, the CSV files, the Jupyter notebooks, and the code that was used to generate the plots are available at https://github.com/COBA-NIH/2023_ImageAnalysisSurvey/tree/main. Interactive versions of most figures are available via Streamlit^21^ at https://broad.io/2022SurveyApp.

