## Supplementary material for "Bridging Imaging Users to Imaging Analysis - A community survey"

**Supplementary figures**

**Supplementary figure 1**

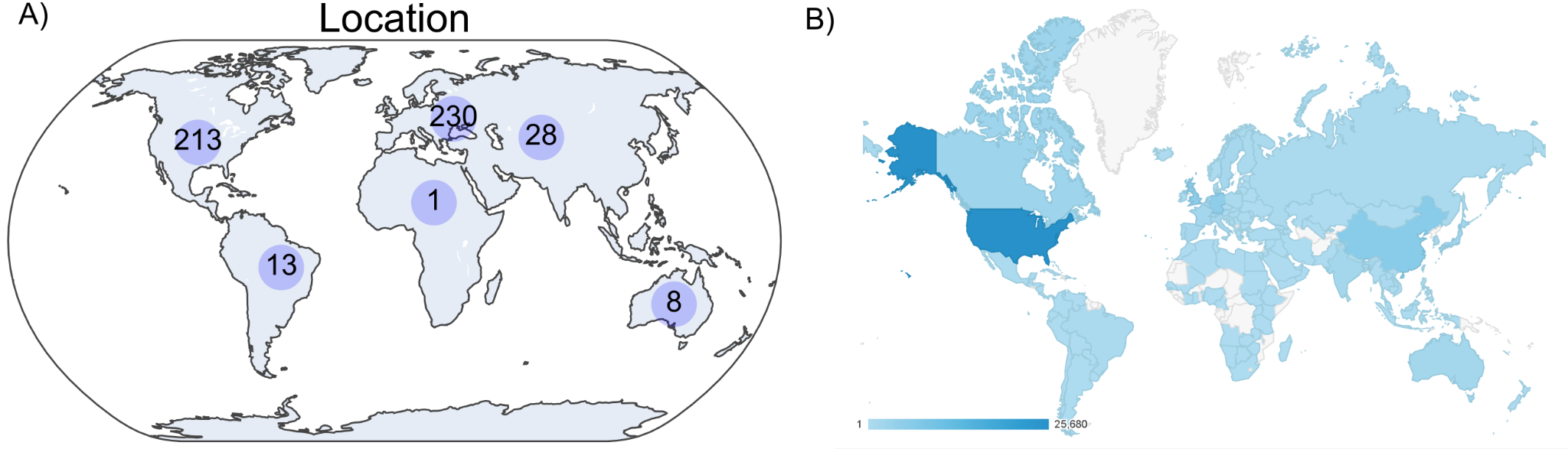

**Figure S1- Survey respondents were drawn primarily from North America and Europe**

A) *Answers to the multiple-choice question “Where do you currently primarily work?”*
B) *Image from Google Analytics showing the number of visitors to the Cellprofiler website in the year 2022. The scale bar indicates the number of visitors.*

**Supplementary figure 2**

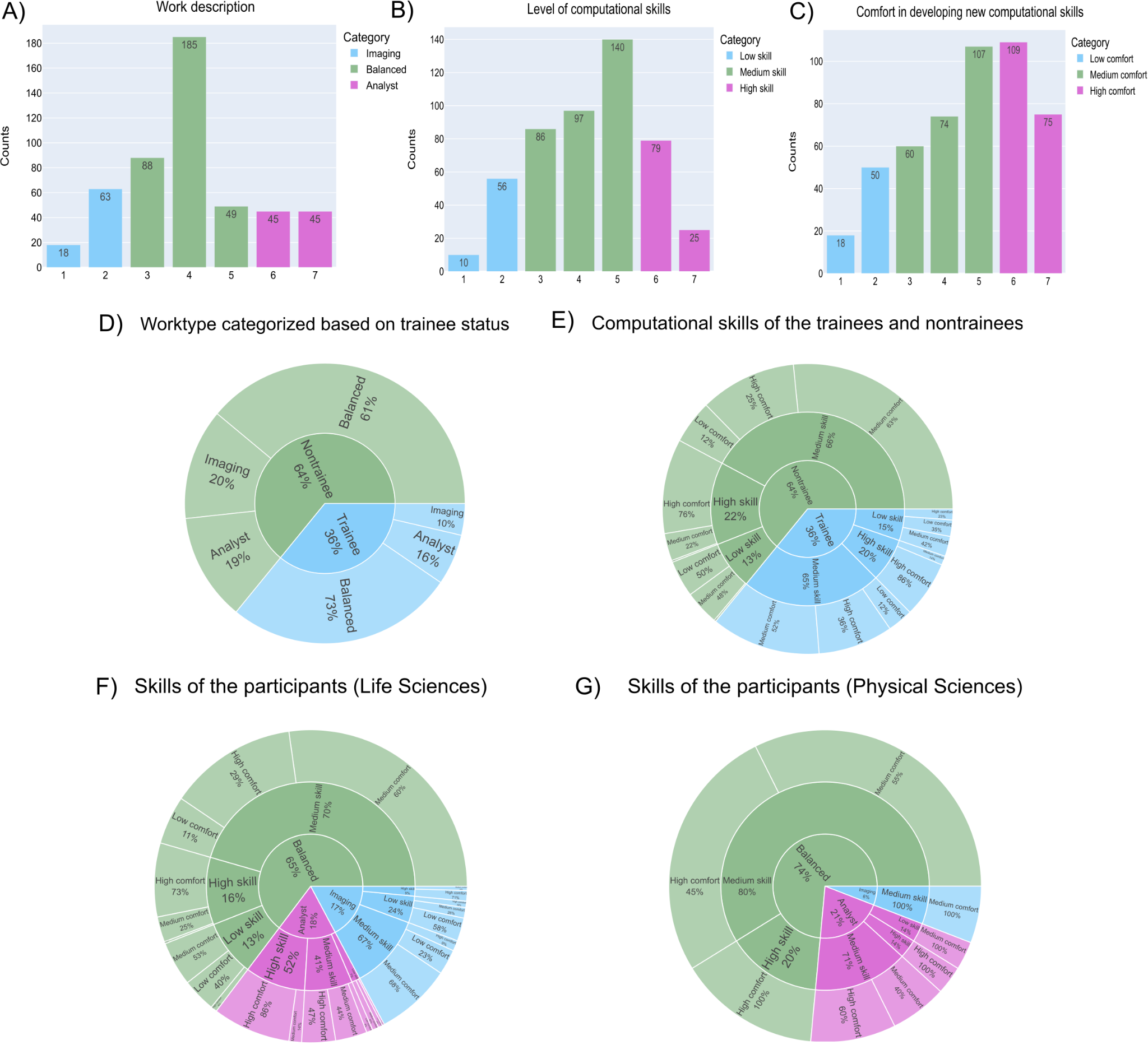

**Figure S2**

**Work description and skills of the participants**

A) *Answers to the question “How would you describe your work?” on a linear scale of 1 to 7 with 1 defined as ‘Nearly entirely imaging (sample prep, optimizing/deciding on imaging modalities, acquiring images and data, etc)’ and 7 defined as ‘Nearly entirely image analysis (finding the right tools to analyze a particular experiment, optimizing the analysis, data mining)’. Scale values 1 and 2 were categorized as “Imaging” work type, values 3 to 5 were categorized as “Balanced” work type, and values 6 and 7 were categorized as “Analyst”.*
B) *Answers to the question “How would you rate your computational skills?” on a linear scale of 1 to 7 with 1 defined as ‘Very poor’ and 7 defined as ‘Excellent’. Scale values 1 and 2 were categorized as “Low skill”, values 3 to 5 were categorized as “Medium skill” and values 6 and 7 were categorized as “High skill”.*
C) *Answers to the question “How would you rate your comfort in developing new computational skills?” on a linear scale of 1 to 7 with 1 defined as “Very uncomfortable” and 7 defined as “Very comfortable”. Scale values 1 and 2 were categorized as “Low comfort”, values 3 to 5 were categorized as “Medium comfort” and values 6 and 7 were categorized as “High comfort”.*
D) *Answers to the question “How would you describe your work?” were categorized as described in part A and the answers - ‘‘Undergraduate/Graduate student’ and ‘Postdoctoral fellow’ to the question “Which of the following roles best describes you?” were considered as ‘Trainees’ and other roles were considered as ‘Nontrainees’.*
E) *Answers to the question “How would you rate your computational skills?” and “How would you rate your comfort in developing new computational skills?” were categorized based on the trainee status.*
F) *Answers to part A, B, and C were classified based on the responses to the question “The next question will ask you about particular image analysis tools and techniques. Do you want to answer questions about microscopy in the field/area of life sciences or physical sciences?” with only ‘Life Sciences’ participants replies represented in the figure.*
G) *Answers to part A, B, and C were classified based on the responses to the question “The next question will ask you about particular image analysis tools and techniques. Do you want to answer questions about microscopy in the field/area of life sciences or physical sciences?” with only Physical Sciences’ participants replies represented in the figure.*

**Supplementary figure 3**

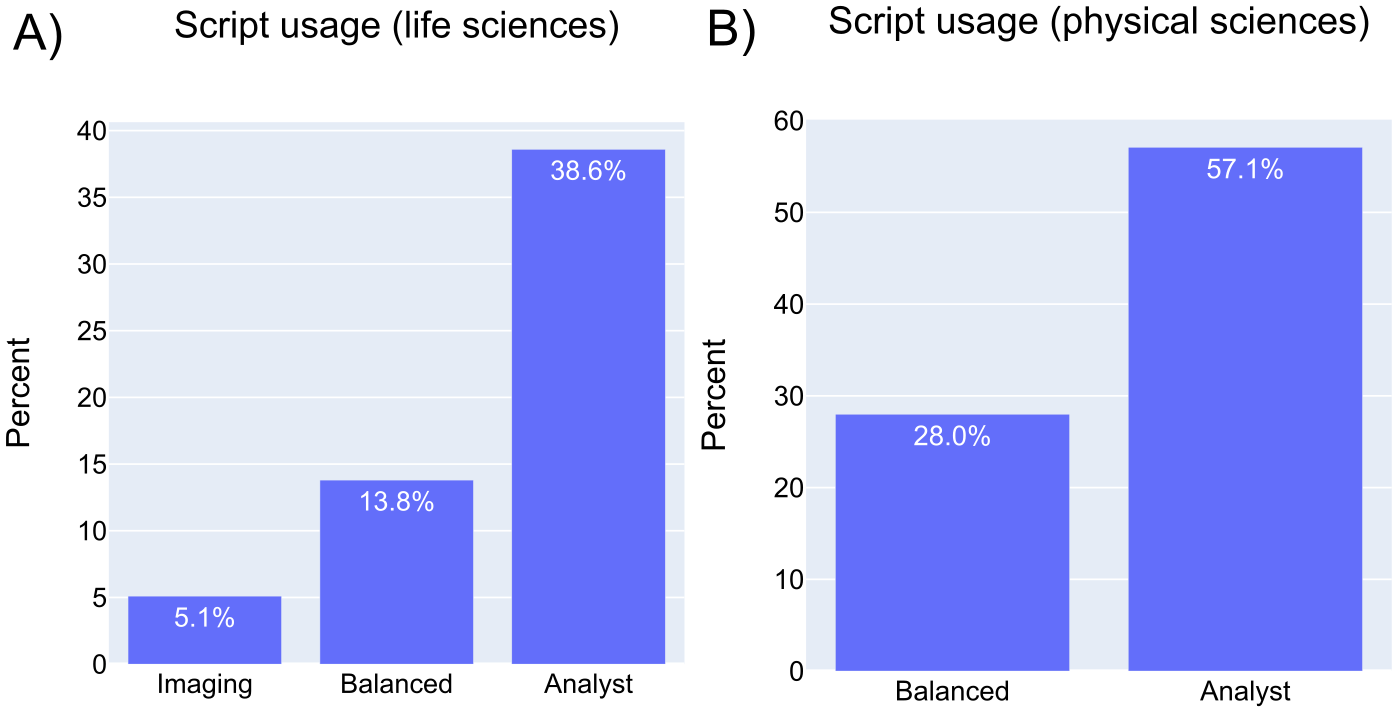

**Figure S3**

**Usage of computational scripts among different work type of the participants**

A) *Usage of computational scripts and libraries among life science participants was categorized based on the ‘work type’ as described in supplementary figure 2.*
B) *Usage of computational scripts and libraries among physical science participants was categorized based on the ‘work type’ as described in supplementary figure 2.*

**Supplementary figure 4**

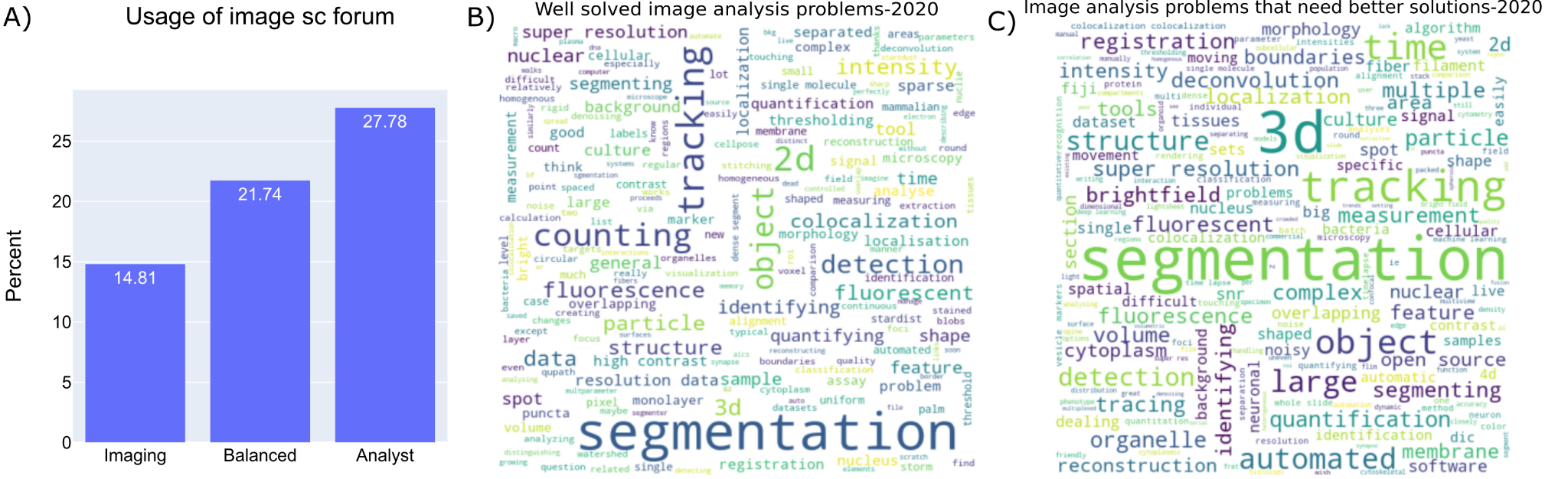

**Figure S4**

**Image analysis problems as stated by 2020 survey participants**

A) *Percentage usage of image sc forum was calculated based on answers provided for the question ‘How do you generally go about solving an image analysis problem? Check the approach(es) you use the most’ normalized with the work type that was categorized based on the answers provided to “How would you describe your work?”. Categorization is given in detail in Figure S2A.*
A) *Wordcloud representation of the unigrams of the answers provided by the 2020 bioimage analysis survey participants to an open-ended question “What image analysis problems (i.e. finding nuclei, tissue analysis, analysis of super-resolution data, etc) do you think are generally well-solved?”.*
B) *Wordcloud representation of the unigrams of the answers provided by the 2020 bioimage analysis survey participants to an open-ended question “What image analysis problems (i.e. finding nuclei, tissue analysis, analysis of super-resolution data, etc) do you wish had easier/better solutions?”.*

**Supplementary figure 5**

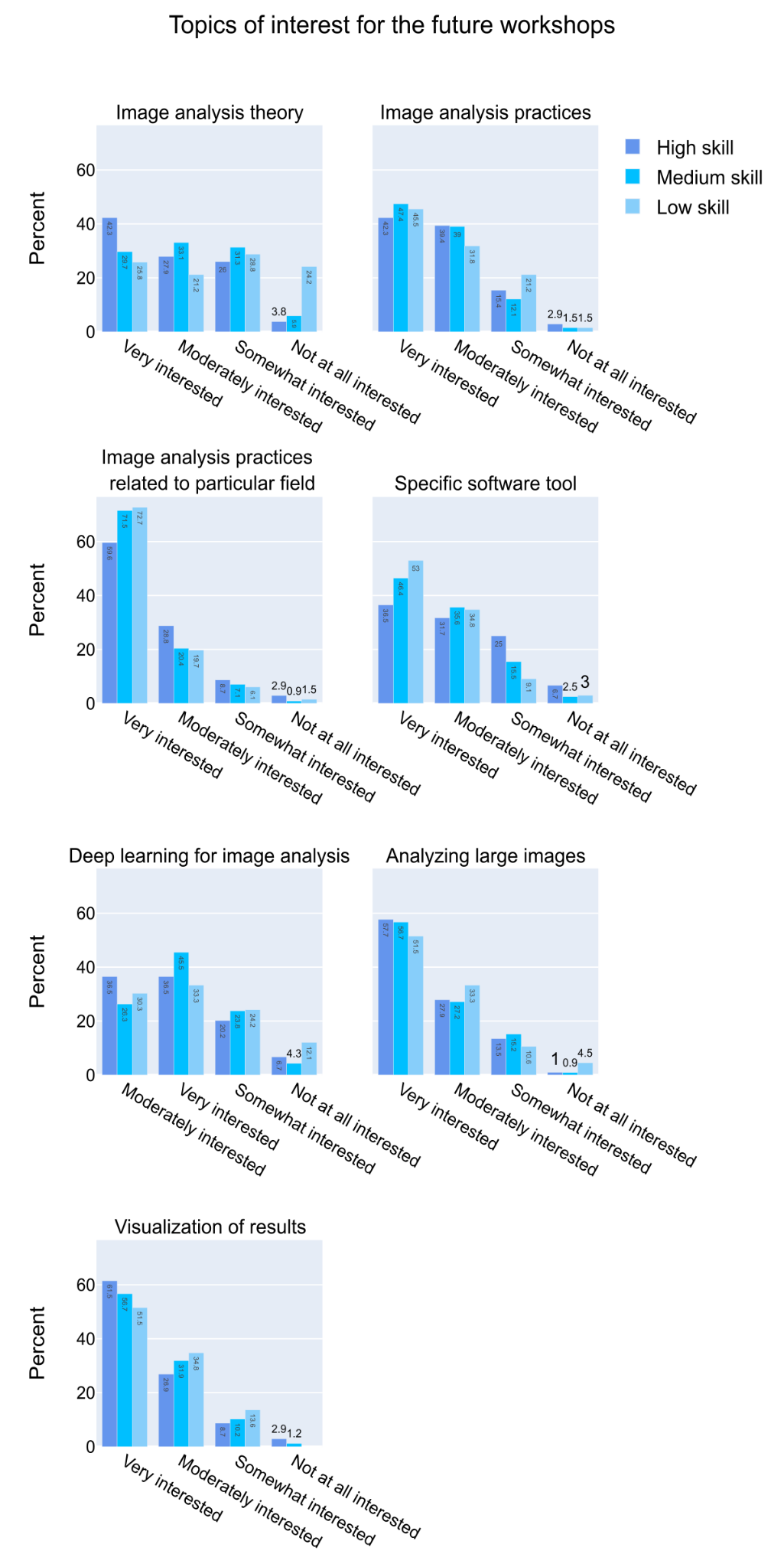

**Figure S5**

**‘Image analysis practices related to sub-discipline’ is the most preferred image analysis topic across all computational skill levels**

*Answers to a multiple-choice grid question “How interested are you in learning more about the following topics?” are categorized based on ‘Level of computational skills’ as described in supplementary S2.*

**Supplementary figure 6**

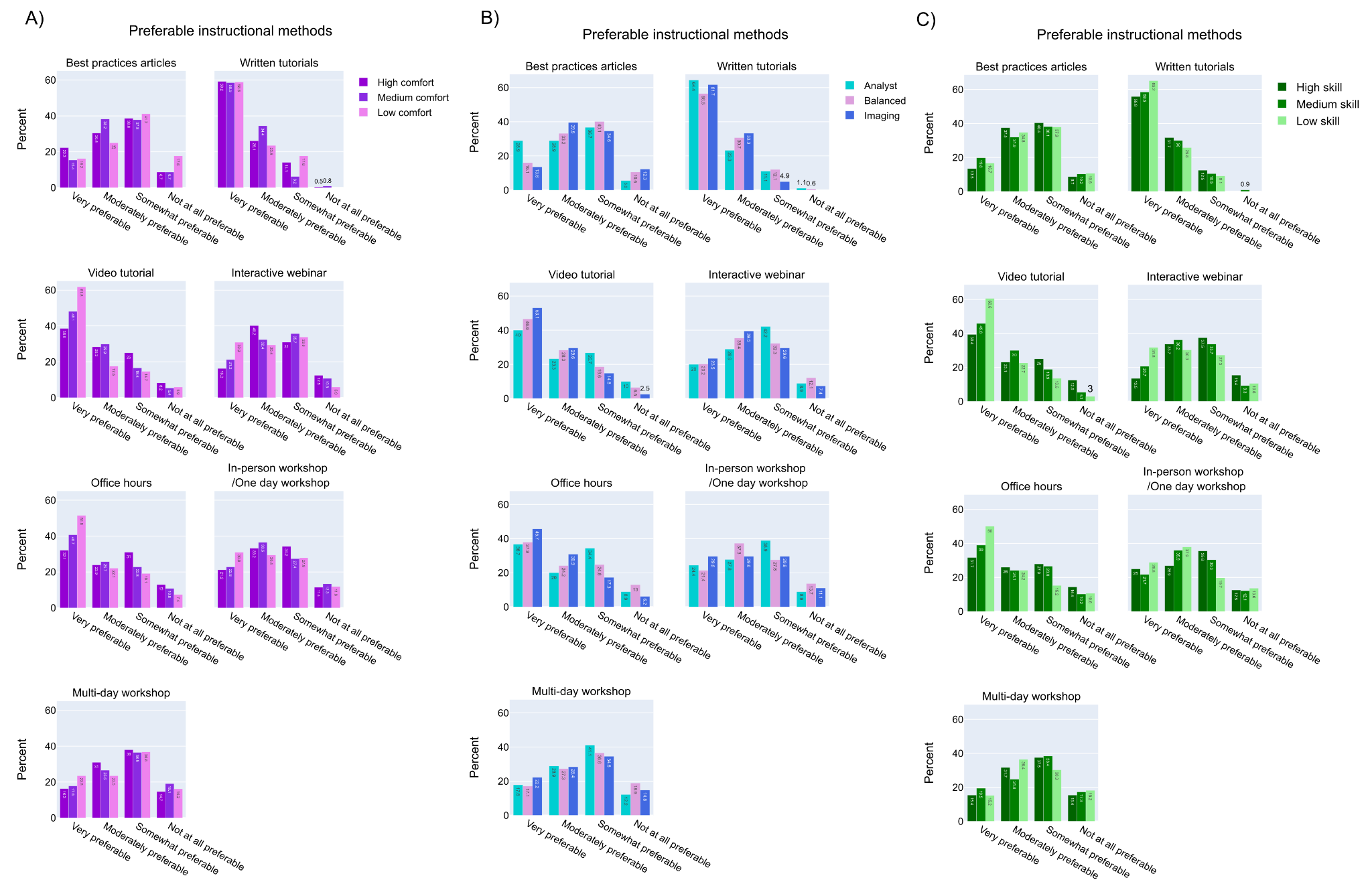

**Figure S6**

**Preference for instructional methods varies by work type, comfort level, and skill level**

A) *Answers to a multiple-choice grid question “In regards to learning more about image analysis, how preferable do you find each of these instructional methods?” are categorized based on ‘Comfort level in developing new computational skills’ as described in supplementary S2C.*
B) *Answers to a multiple-choice grid question “In regards to learning more about image analysis, how preferable do you find each of these instructional methods?” are categorized based on ‘work type’ as described in supplementary S2A.*
C) *Answers to a multiple-choice grid question “In regards to learning more about image analysis, how preferable do you find each of these instructional methods?” are categorized based on ‘Level of computational skills’ as described in supplementary S2B.*

**Supplementary figure 7**

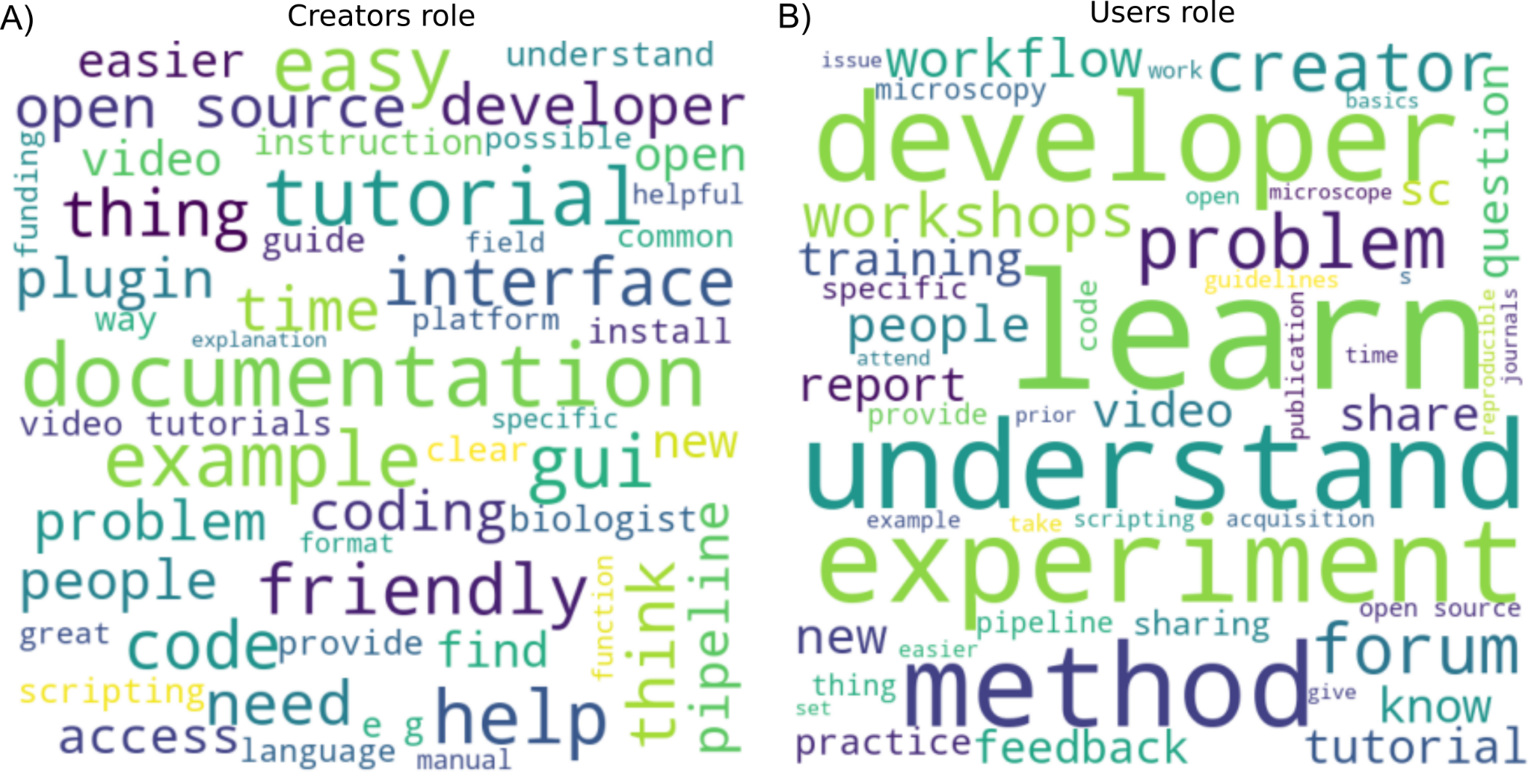

**Figure S7**

**Creators and users role as suggested by the participants**

A) *Wordcloud representation of the unigrams of the answers to an open-ended question “What do you think analysis tool CREATORS (such as software developers)could/should do to make image analysis better and more successful? How best could we encourage them to do it?”*
B) *Wordcloud representation of the unigrams of the answers to an open-ended question “What do you think analysis tool USERS (such as microscopists) could/should do to make image analysis better and more successful? How best could we encourage them to do it?”*

**Supplementary figure 8**

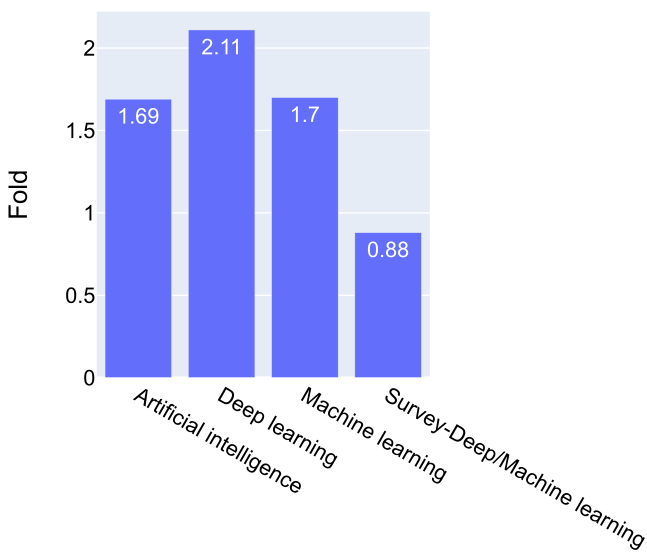

**Figure S8**

**Interest in machine learning/deep learning**

*Fold change in the number of articles that were published in PubMed with the terms - ‘Artificial Intelligence’, ‘Machine learning’, ‘Deep learning’ in 2020 and 2022 were plotted along with the interest level in ‘Deep learning as applied to image analysis’ as shown in the Figure 7C and described in the methods section. A fold change of less than 1 represents decreased interest.*
